# A Genomic View of Environmental and Life History Controls on Microbial Nitrogen Acquisition Strategies

**DOI:** 10.1101/2023.08.10.552805

**Authors:** Linta Reji, Romain Darnajoux, Xinning Zhang

**Affiliations:** Department of Geosciences, Princeton University, Princeton NJ; High Meadows Environmental Institute, Princeton University, Princeton NJ

## Abstract

Microorganisms have evolved diverse strategies to acquire the vital element nitrogen (N) from the environment. Ecological and physiological controls on the distribution of these strategies among microbes remain unclear. Here we examine the distribution of 10 major N-acquisition strategies in taxonomically and metabolically diverse microbial genomes, including those from the Genomic Catalog of Earth’s Microbiomes dataset. We utilize a marker gene-based approach to assess relationships between N acquisition strategy prevalence and microbial life history strategies. Our results underscore energetic costs of assimilation as a broad control on strategy distribution. The most prevalent strategies are the uptake of ammonium and simple amino acids, while biological nitrogen fixation is the least common. Deviations from this energy-based framework include the higher-than-expected prevalence of the assimilatory pathway for chitin, a large organic polymer. Notably, oxygen-respiring chemoorganotrophic and phototrophic microbes possess ∼2-fold higher numbers of total strategies compared to anaerobic microbes. Environmental controls on N acquisition are evidenced by the enrichment of inorganic N assimilation strategies among free-living taxa compared to host-associated taxa. Physiological constrains such as pathway incompatibility add further complexity to N-acquisition strategy distributions. Finally, we discuss the necessity for microbially-relevant environmental metadata for improving mechanistic and prediction-oriented analyses of genomic data.

## INTRODUCTION

Nitrogen (N) is a critical element for life, and microorganisms have evolved diverse strategies for acquiring N from the environment. These N acquisition strategies target various N forms, including the inorganic forms dinitrogen gas, ammonia/ammonium, nitrite, and nitrate; nitrogenous organic compounds such as urea, cyanate, and amino acids; as well as large N-containing polymers such as chitin and peptidoglycan. The energetic costs of uptake and assimilation (i.e., ATP, reducing equivalents) vary across compound classes, and this is thought to regulate N substrate preferences among microbes (1). Various lines of ample evidence indicate that microbes downregulate energetically expensive N assimilation pathways such as biological nitrogen fixation or chitin depolymerization when small, more cheaply assimilated fixed N compounds are available (e.g., 2–5). Whether microbial genomic architectures reflect these differences in substrate preferences remains unclear.

Cellular costs of uptake and assimilation vary depending on the molecular size and redox state of the N compound. N assimilation in most microbes proceeds via the incorporation of ammonium into carbon skeletons through the glutamine synthase-glutamate synthase (GS-GOGAT) cycle, producing glutamine and glutamate, which are incorporated into cellular anabolism. While the uncharged species ammonia (NH_3_) can enter the cell via diffusion when it is abundant in the environment (e.g., alkaline conditions; 6), specialized transporter proteins are widely used by microbes to transport both NH_3_ and the protonated form ammonium (NH_4_^+^) into the cell (7). The GS/GOGAT pathway consumes 1 ATP per molecule of NH_4_^+^.

Amino acids are another major class of low molecular weight (LMW) organic N compounds available for microbes in the environment. While amino acids are expected to be assimilated relatively easily to cellular macromolecules, their uptake can involve energetic investment in the form of proton/sodium ion motive force (PMF/SMF) and/or ATP (8). Most amino acid transporters are PMF or SMF-dependent secondary active transporters (8) (i.e., move substrates against the concentration gradient while transporting other solutes down their concentration gradient). As amino acids can also serve as a carbon and sulfur source, microbial uptake of amino acids does not necessarily reflect their nitrogen strategy. In terms of energetic investment, amino acid assimilation should be equally if not more favorable compared to ammonium given the minimal post-uptake modifications required. Experimental evidence indicates that dissolved amino acids can repress ammonium uptake and assimilation, at least in heterotrophic microbes (9,10).

Assimilating N sources other than NH_3_/NH_4_^+^ or amino acids incurs additional energetic investments in terms of specialized transporters for uptake as well as intracellular reduction of the substrate to NH_4_^+^. For example, assimilating one molecule each of nitrite (NO_2_^-^) and nitrate (NO_3_^-^) consumes 6 and 8 reducing equivalents each, respectively. In comparison, biological nitrogen fixation (BNF), the process by which microbes assimilate atmospheric N_2_ requires 8 reducing equivalents and 16 ATPs per mole of N_2_ fixed. Additional indirect costs for assimilation include making and maintaining the complex enzymatic machinery required for each process, particularly BNF. Acquiring high molecular weight (HMW) organic N sources also require significant metabolic energy input due to the diverse and bulky nature of these compounds. Microbes must not only synthesize a diverse array of transporters to target different HMW compounds but also need to produce various extracellular enzymes to break down large polymers in the extracellular environment prior to uptake.

Based on the relative difficulty with which microbes can access diverse N sources, Norman and Freisen (1) proposed a N acquisition strategy for free-living N_2_-fixing microbes (diazotrophs) under fixed N limitation, which postulates a preference for low molecular weight (LMW) organic N over BNF, followed by high-molecular weight (HMW) organic N. Accounting for the relative energy investments required for accessing, transporting, and assimilating different N sources, this ‘LAH strategy’ (which stands for ‘LMW N – atmospheric N_2_ – HMW N’) proposes the following order of N compound preferences for free-living diazotrophs: ammonia/ammonium > nitrite/nitrate > LMW organic N > BNF > HMW organic N. Norman and Freisen (1) indeed found evidence from pure culture growth experiments and genomes that several lineages of free-living soil diazotrophs can access the HMW-N pool (primarily via extracellular protein degradation). Under the LAH framework and the differential energetic investments involved in the assimilation of various N sources, it may be presumed that the relative distribution of N acquisition strategies among microorganisms follow their respective substrate preferences.

The broader validity of the LAH strategy remains to be tested, however, as the realized cost of N acquisition likely varies depending on a microbe’s environmental context and life history adaptations (i.e., mode of metabolism, resource allocation, motility, etc.). For instance, the dynamic nature of the bioavailable N pool as well as spatial heterogeneity of geochemistry in structured environments such as soils (11) may shape the relative distributions of N acquisition strategies within natural microbial communities. Several outstanding questions in this regard are: (i) how prevalent are different N acquisition strategies among microbes across environmental systems?, (ii) are specific pathway combinations preferred over others?, (iii) how do life history traits (e.g., mode of metabolism, resource allocation, motility, etc.) and environmental context affect the number of N-acquisition pathways present in an organism?

To address these questions, we conducted a genomic study examining the distribution of ten major N acquisition strategies across microbial genomes with taxonomic, metabolic and habitat diversity. We utilized metagenome-assembled genomes (MAGs) from the Genomic Catalog of Earth’s microbiomes (GEMs) dataset, which consists of >10,000 metagenomes representing diverse environmental systems (12). We also conducted a targeted analysis of 6 microbial clades chosen based on diverging metabolic and evolutionary relatedness, including complete genomes of the phylum *Cyanobacteria*, genus Clostridium, and the orders *Rhizobiales, Desulfovibrionales, Methanobacterales*, and *Methanococcales*. The presence both diazotrophic and non-diazotrophic lineages within each clade enabled more comprehensive examination of N acquisition capabilities. In each genome set, we employed a marker gene-based approach to systematically catalogue the prevalence of the following 10 major N acquisition strategies: biological nitrogen fixation (BNF), ammonium uptake, ferredoxin-dependent nitrite reduction; NADH-dependent nitrite reduction; assimilatory nitrate reduction; urea uptake; cyanate uptake; chitin depolymerization; assimilation of chitin oligomers (i.e., beta-N-acetyl hexosamine); and amino acid uptake. We then examined the influence of metabolic adaptations and environmental context on the distribution of these N acquisition strategies among diverse groups of microbes.

## MATERIALS AND METHODS

### Compiling and pre-processing genomes

Genome assemblies from the Genomic Catalog of Earth’s microbiomes (GEMs) dataset were downloaded from https://portal.nersc.gov/GEM/ (12). The original set of 52,515 MAGs was filtered to only retain those meeting the MIMAG high quality genome criteria (13). This resulted in 9,143 MAGs after quality filtering. In addition to the GEMs set, complete genome sequences of organisms belonging to five selected clades – *Cyanobacteria, Clostridium, Rhizobiales, Desulfovibrionales, Methanobacterales*, and *Methanococcales* – were downloaded from the NCBI RefSeq database using genome_updater (https://github.com/pirovc/genome_updater) in March 2022. All genome assemblies were analyzed using Prodigal (14) to predict protein-coding sequences.

### Examining N-acquisition strategy prevalence across genomes

We examined the incidence of ten N-acquisition strategies using specific functional genes indicative of each strategy. Marker gene-based surveys can yield false positive results if genes encoding critical enzyme subunits or structural components are absent from the genome. To mitigate such artifacts, we focused on functionally critical elements of the key enzyme associated with each pathway. The following pathway-marker gene combinations were used in our analysis: ammonium uptake (marker gene: ammonium permease, AmtB); biological nitrogen fixation (nitrogenase, NifHDK); Ferredoxin (Fd)-dependent nitrite assimilation (Fd-nitrite reductase, NirA); NADH-dependent nitrite assimilation (NADH-nitrite reductase, NirBD); nitrate assimilation (assimilatory nitrate reductase, NasA co-occurring with an assimilatory nitrite reductase); cyanate assimilation (cyanase, CynS); urea assimilation, (urease, UreC), Chitin depolymerization (chitinase), N-acetyl hexosamine assimilation (N-acetyl hexosaminidase), and amino acid uptake (various amino acid transporters).

For strategies other than amino acid uptake, reference protein sequences for each pathway marker were obtained from the NCBI RefSeq protein database (Table S1). Reference sequences belonging to five distinct families of amino acids (all within the APC superfamily) were obtained from the Transporter Classification Database (TCDB; ref. 15,16; Table S1). For each marker protein, multiple sequence alignments (MSAs) were generated using MAFFT (v7.475; ref. 17). These alignments were then used to generate profile HMMs for each protein via hmmbuild in HMMER (http://hmmer.org/). Following this, hmmsearch was used to identify homologs of each marker protein across all genomes. The E-value thresholds for hit identification were adjusted based on BLASTp search results against the UNIPROT/SWISSPORT databases (18).

The putative hits for each marker protein were then aligned with reference sequences using MAFFT (v7.475; ref. 17), and individual phylogenetic trees were generated using FastTree (v2.1.10; ref.19,20). The trees were inspected in Taxonium (21) and manually curated. Curation involved BLASTp searches of sequences from each major phylogenetic cluster against the RefSeq as well as the Uniprot/Swissport databases. Putative amino acid transporter hits were searched against the TCDB (15,16) to assess homology. False positives identified based on homology searches were removed from the final list of hits for each marker. Curation was particularly challenging for nitrate and nitrite reductases as the hits often included diverse kinds of oxidoreductases. Erring on the side of caution, we discarded all sequences that could not be confidently classified as a NasA or NirA. Curated results are summarized in Table S2.

### Statistical analyses

All statistical analyses were carried out in R (v4.1.2). Of the 9,143 genomes in the filtered GEMs dataset, one did not have a Domain-level taxonomic identification. Additionally, 274 genomes did not contain homologs of any of the ten examined pathways. Those missing any of the 10 examined pathways were not analyzed further as it was impossible to determine if this was an artifact of genome incompleteness. The final filtered GEMs set included 8,868 genomes, consisting of 8,696 bacteria and 172 archaea.

For comparisons involving metabolic adaptations, we classified the genomes as “aerobic”, “facultatively anaerobic”, and “anaerobic”. The genomes were also classified as “photoautotrophic”, “photoheterotrophic”, “chemoautotrophic”, “chemoorganotrophic”, and “mixotrophic”. These codings were assigned based on expected metabolic strategy within a conserved taxonomic level based on what has been reported in the literature (for example, all genomes within class *Cyanobacteriia* were coded as aerobic photoautotrophs). Lineages lacking physiologically characterized members were excluded from the GEMs dataset, which resulted in a final set of 4,826 genomes for metabolism-based analyses (Table S3).

Differences in N strategy incidence/prevalence distributions across various metadata variable groupings were statistically tested using Kruskal-Wallis tests followed by post-hoc pairwise comparisons using the Wilcoxon Rank Sum test. Generalized mixed effects linear models were used to compare associations between various N-acquisition strategy metrics and metadata variables, controlling for taxonomic relatedness between genomes. Dimensionality reduction was carried out by using prcomp (https://www.rdocumentation.org/packages/stats/versions/3.6.2/topics/prcomp) and logisticPCA (22).

## RESULTS AND DISCUSSION

### Genomic patterns of microbial N acquisition indicate importance of ammonium and organic N compounds

Based upon the expected energetic costs of uptake and assimilation, as proposed under the LAH framework, microbial N acquisition is expected to follow a predictable order of compound preferences (1). More easily assimilated compounds such as inorganic N (ammonium, nitrite, and nitrate) and low-molecular weight organic N (amino acids, urea, cyanate) compounds are preferred over BNF and high-molecular weight organic N compounds. Subsequently, it can be inferred that the incidence of the various N acquisition strategies in microbial genomes tracks the relative order of N compound preferences. For example, inorganic and LMW organic N uptake pathways would be more commonly found among microbes compared to BNF and HMW-N uptake pathways. In order to test this prediction, we utilized the recently published GEMs dataset, which includes 52,515 genomes assembled from 10,450 metagenomes originating from diverse environments (12).

In examining the GEMs genomes, amino acid uptake emerges as the most prevalent among the ten N strategies examined, closely followed by ammonium uptake (Fig. 1a). This likely reflects the fact that ammonium and amino acid incorporation necessitates minimal downstream modification of the acquired substrate. Some forms of amino acids may be preferred over others, however. Our analysis included 5 subfamilies of amino acid transporters as defined within the TCDB (15,16): betaine/carnitine/choline (BCCT) family, alanine/glycine:cation symporter (AGCS) family, branched chain amino acid:cation symporter (LIVCS) family; hydroxy/aromatic amino acid permease (HAAAP) family; and the APC family consisting of those not falling within any of the remaining four families. Among these, BCCT, LIVCS, and HAAAP families target more complex amino acid forms compared to AGCS. Indeed, AGCS transporters were significantly more prevalent than branched-chain/aromatic amino acid transporters (Fig. 1b). Thus, a key determinant of N-substrate preference might be the extent of downstream modification of the substrate before incorporation into anabolism.

**Figure 1:**
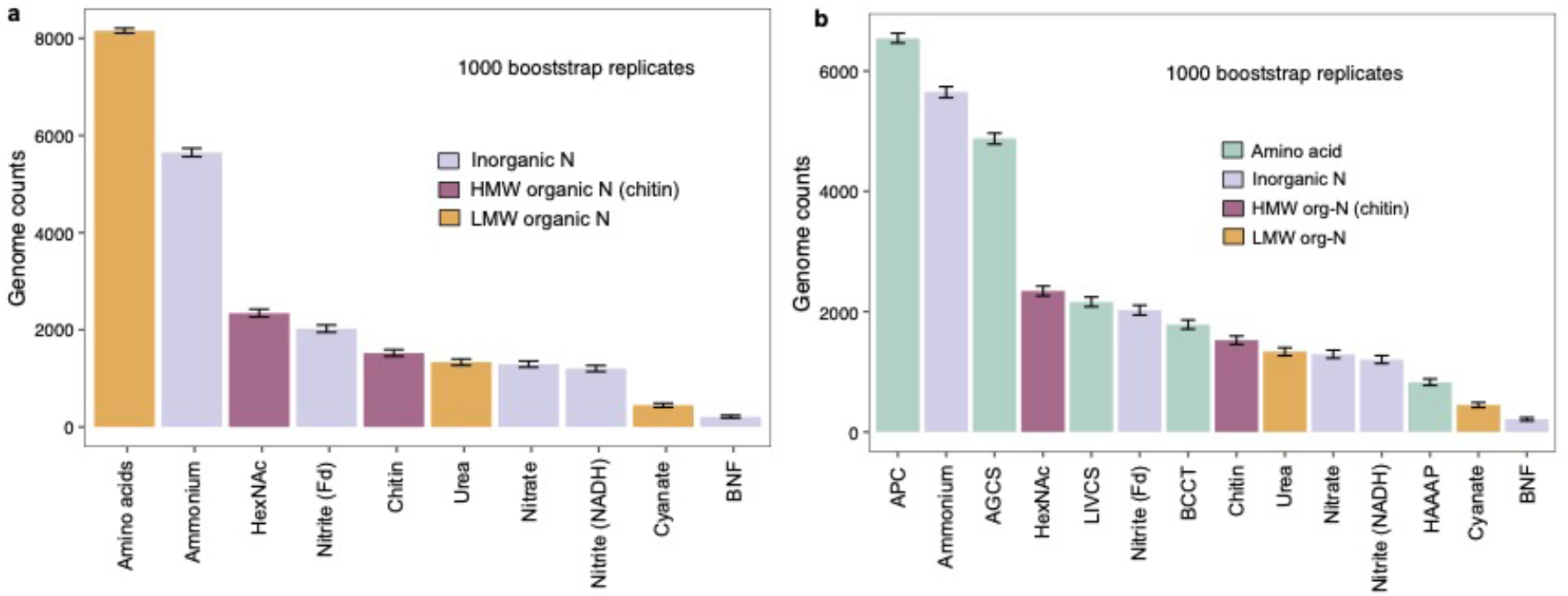
Amino acid and ammonium acquisition dominate over all other N acquisition strategies in microbial genomes; and High Molecular Weight (HMW)-organic N assimilation may be more common than inorganic N acquisition. a. Prevalence of the various N-compound acquisition strategies among genomes in the filtered GEMs dataset. HexNAc: N-acetyl-hexosamine, a compound class that includes chitin monomers; BNF: biological nitrogen fixation. A non-parametric bootstrapping approach (1000 bootstrap replicates) was used to reduce biases from uneven sampling depths. Bars indicate the bootstrapped median number of genomes containing each strategy; error bars indicate the 95% bootstrap confidence intervals around the median. **b**. Prevalence patterns presented in panel a, with greater resolution into the relative prevalence of amino acid transporter families. AGCS, alanine/glycine:cation symporter; LIVCS, branched chain amino acid:cation symporter; BCCT, betaine/carnitine/choline transporter; HAAAP, hydroxy/aromatic amino acid permease; APC, Amino Acid-Polyamine-Organocation family. The latter (APC) consists of amino acid transporters not included within the remaining 4 families.

Another notable result from the GEMs analysis is the higher-than expected prevalence of genes involved in chitin depolymerization and the uptake of chitin monomers (i.e., N-acetyl-hexosamine, HexNAc). Chitinases and N-acetyl-hexosaminidases are more common compared to nitrite, nitrate, urea, and cyanate assimilation genes (Fig. 1). These patterns were robust to multiple resampling approaches performed to reduce taxonomic sampling bias (Fig. S1). However, we do not fully discard biases resulting from overrepresentation of certain habitat types (e.g., human and other host microbiomes) in the GEMs dataset.

Chitin-related genes were more prevalent than genes involved in the assimilation of smaller organic N compounds such as urea and cyanate (Fig. 1, S1). The prevalence of chitinases might be due to the widespread availability of chitin as it is the second most abundant biopolymer in the environment (23). Yet it remains unclear why chitin degradation may be more prevalent than urea assimilation. This could be related to the relative stability of each compound in the environment (i.e., chitin generally more stable than urea, which readily dissociates into ammonia and carbon dioxide). In the case of cyanate, however, the relatively low availability of this compound in the environment (up to ∼10,000-fold lower in availability compared to ammonium across habitat types (24)) may be a contributing factor.

Among the inorganic-N strategies, excluding ammonium uptake, ferredoxin-dependent nitrite reduction appears to be the dominant strategy (Fig. 1 and S1). As would be predicted based upon the energetic cost of assimilation paradigm, BNF is the least prevalent strategy among the GEMs genomes. Notably, however, this contrasts with the predictions of the LAH framework, which suggested that atmospheric N_2_ assimilation might be preferred over HMW-N acquisition.

Hence, the genomic patterns observed in the GEMs dataset (Fig. 1) indicate a slightly different order of N-compound preferences compared to the predictions of the LAH framework (Fig. 2). Simple amino acids may be equally if not more preferred over ammonium given similar energetic investments in the uptake/assimilation of these compounds. HMW organic-N assimilation may be more prevalent than recognized, compared to specialized strategies targeting LMW organic compounds such as urea and cyanate. As these patterns indicate, energetics of uptake/assimilation do not sufficiently explain the relative distributions of the different strategies, suggesting more complex environmental and life history controls on pathway distributions. We examine these next.

**Figure 2:**
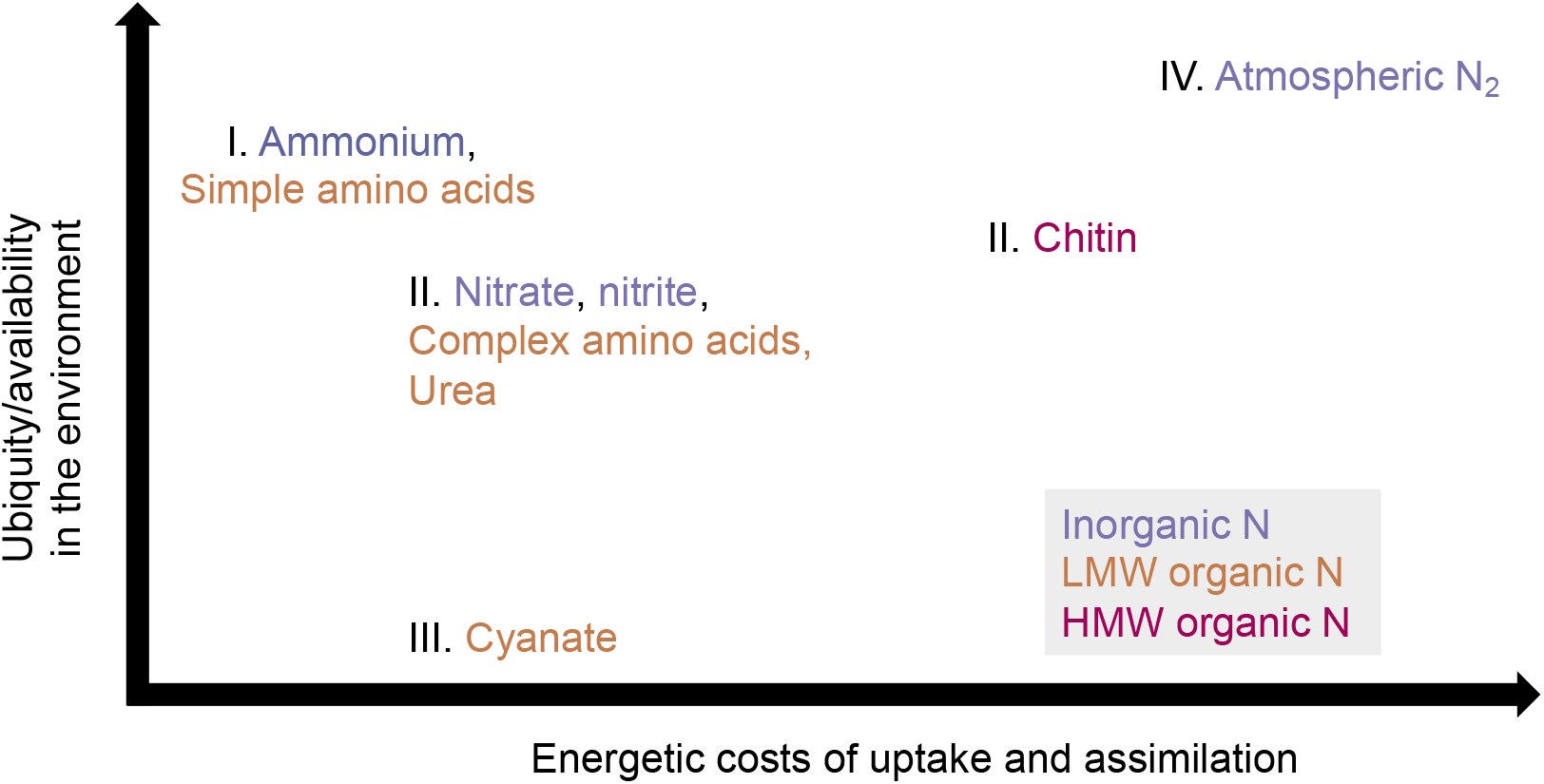
Microbial N-compound preferences indicated by the genomic analysis. The order of compound preferences likely follows the energetic costs of assimilation as well as relative compound availabilities, as discussed in the text. The Roman numerals indicate inferred order of compound preferences based on the GEMs analysis. Additional controls on the incidence of N strategies not depicted here includes physiological factors such as metabolic toxicity (i.e., oxygen toxicity of BNF) and pathway incompatibility (e.g., nitrite interfering with sulfite reduction in sulfate reducers).

### The abundance of microbial N acquisition strategies scale with genome size and metabolic energy yield

Given the differential direct and indirect energetic and cellular costs of N acquisition strategies, we hypothesized that the distribution and diversity of the different strategies would vary as a function of metabolic energy yield. Metabolic modes yielding higher free energy per reaction (e.g., oxygen respiration) likely result in higher cellular N demand (e.g., due to faster growth rate and associated faster turnover of cellular components) while at the same time affording the microbe the ability to maintain and use multiple uptake/assimilation pathways. In testing this hypothesis, we considered two fundamental aspects of the metabolic strategy of an organism: mode of respiration (or the lack thereof; coded as anerobic, aerobic, or facultative anerobic in our analyses) and energy metabolism (chemoorganotroph, chemoautotroph, photoautotroph, and mixotroph). First, we assessed if larger genomes correspond with higher number of total N acquisition strategies. We then examined if the number of N acquisition strategies varied across different modes of metabolism.

In analyzing the N strategy distributions within a set of six microbial clades (selected based on the presence of diazotrophic and non-diazotrophic members within each clade, and the diversity of central metabolic strategies) – *Cyanobacteria, Clostridia, Desulfovibrionales, Rhizobiales, Methanobacteriales, Methanococcales* – we observed that (i) the number of N assimilation pathways scaled with genome size, and (ii) lineages characterized by higher metabolic energy yield generally harbored a larger number of N acquisition strategies (Fig. 3a). For example, predominantly aerobic organisms (*Cyanobacteria* and *Rhizobiales*) generally possess higher number of N-acquisition strategies than obligate anaerobes (*Clostridia, Desulfovibrionales*, and methanogens; Fig. 3a). This stands despite the typical habitat overlap between *Rhizobiales* and the non-cyanobacterial clades, suggesting that any effect of physicochemical variability such as differential substrate diffusion and organic matter complexity may have a relatively weaker influence over N assimilation strategy distributions than the primary mode of metabolism. Additionally, the number of N assimilation strategies scale with the genome size among these clades (Fig. 3a), suggesting that aerobic organisms with larger genomes generally have the potential to obtain N from a larger diversity of sources. Conversely, the ability to assimilate a larger diversity of N compounds also likely presents an opportunity to build larger genomes, establishing a reinforcing mechanism.

**Figure 3:**
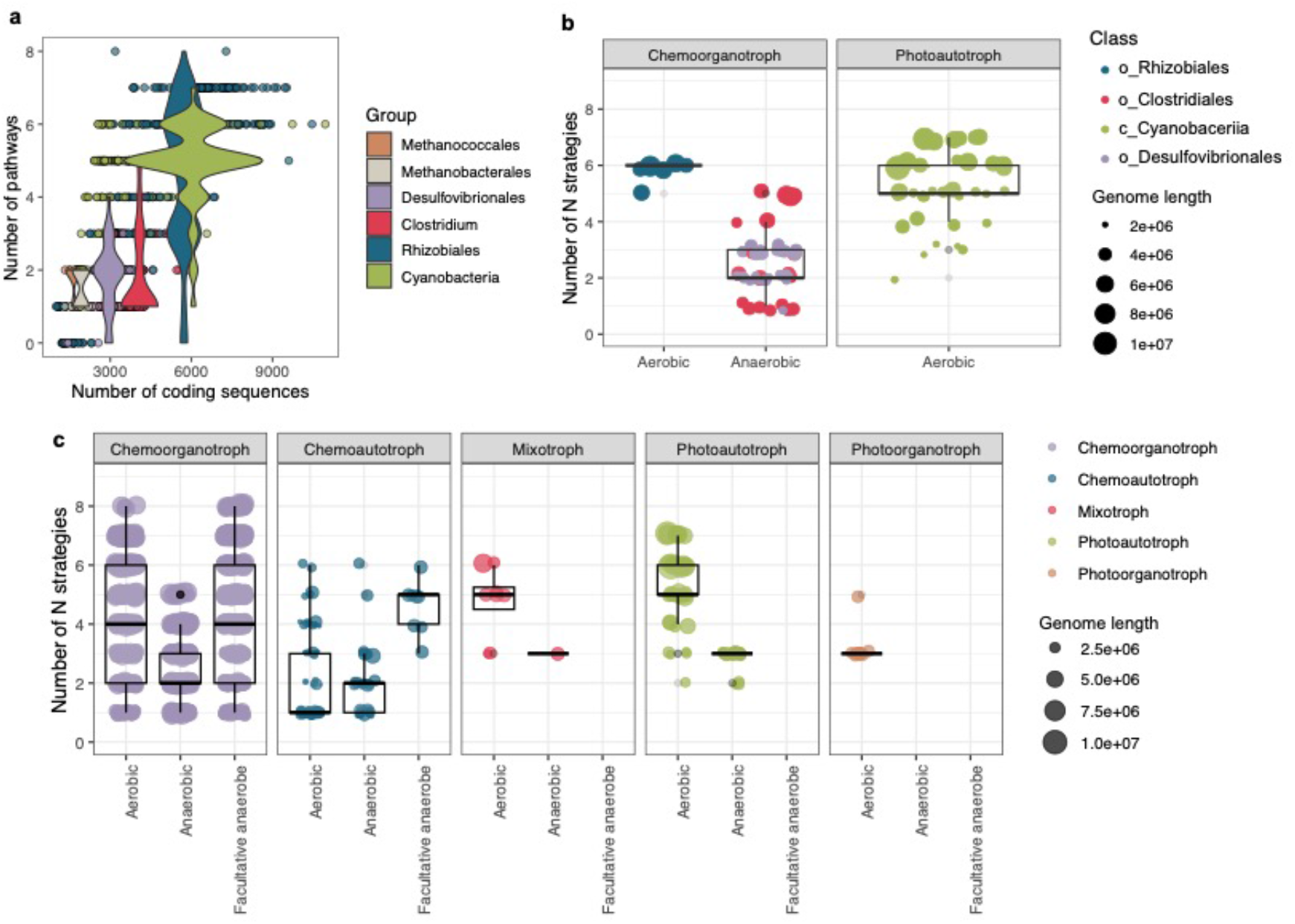
Diversity of N-acquisition strategies scales with microbial genome size and correlates with aerobic respiration. a. The number of N-acquisition pathways detected versus the number of coding sequences in selected phylogenetic clades. The violin plots indicate the distribution of N-pathway counts across clades. **b**. Total number of N-acquisition pathways in the GEMs dataset corresponding to the selected clades presented in panel a, partitioned by metabolic mode. The filtered GEMs set did not include any methanogens. **c**. Total number of N-acquisition pathways across all genomes in the filtered GEMs set (n=4749) partitioned by metabolic mode.

Expanding the analysis to the GEMs, we first examined genomes corresponding to the selected clades above within this larger set of MAGs. The GEMs set, after quality control (See Methods), did not include methanogenic archaeal genomes. Consistent with the patterns observed for the selected clades above, we observed a larger number of total N acquisition pathways in aerobic lineages with larger genomes (Fig. 3b). Observations for the overall filtered GEMs set (4749 genomes) align with these patterns as aerobic and facultatively aerobic organisms possess significantly more N acquisition strategies than anaerobic organisms (Fig. 3c). Mode of metabolic energy generation appears to be an additional control on this as the diversity of acquisition strategies was significantly higher for (a) aerobic phototrophs over anaerobic phototrophs, and (b) facultatively anaerobic chemoautotrophs over aerobic and obligately anaerobic chemoautotrophs (Fig. 3c). In contrast, obligately anaerobic chemoorganotrophs have significantly lower diversity of N acquisition strategies compared to aerobic and facultatively anaerobic chemoorganotrophs (Fig. 3c).

These results also indicate a significant correlation between genome length and the diversity of N acquisition strategies (linear regression on log (genome length) and the number of pathways; p-value <2e-16; Fig. 3c, S2), as seen in the case of the 6 selected clades (Fig. 3a). In addition, there is a significant positive correlation between genome length and aerobic or facultatively anaerobic metabolisms (p-values <2e-16 and 1.82e-05, respectively). In contrast, anerobic respiration is significantly negatively correlated with genome length (p-value <2e-16). It then follows that oxygen-respiring organisms, which tend to have larger genomes, harbor more diverse (larger number of) N acquisition strategies. While these results are expected at the broad scale of metabolic pathway distributions (e.g., larger genomes can accommodate higher number of metabolic pathways), we confirm this with respect to the acquisition of the key nutrient N.

Given the correlation between N pathway counts and metabolic energy yield (Fig. 3), we can expect that higher cost N acquisition strategies will also correlate to metabolic energy yield. To test this hypothesis, we examined whether higher-cost N acquisition pathways (i.e., BNF and chitin acquisition) are more prevalent among oxygen-respiring organisms. Chitinase distribution was significantly positively associated with aerobic respiration (logistic regression, p-value 1.67e-10; Fig. 4a) and facultatively anaerobic respiration (0.04), and negatively with anaerobic respiration (p-value <2e-16). There may be some influence from sampling biases as when controlled for taxonomic relatedness between genomes at the family level using a mixed effects generalized linear model, the positive association between chitinase incidence and aerobic respiration was not significant while it still significantly negatively correlated with anerobic respiration (binomial GLMM; p-value 0.01). Indeed, the majority of chitinase-positive genomes originated from aerobic and facultatively aerobic lineages (Fig. 4a).

**Figure 4:**
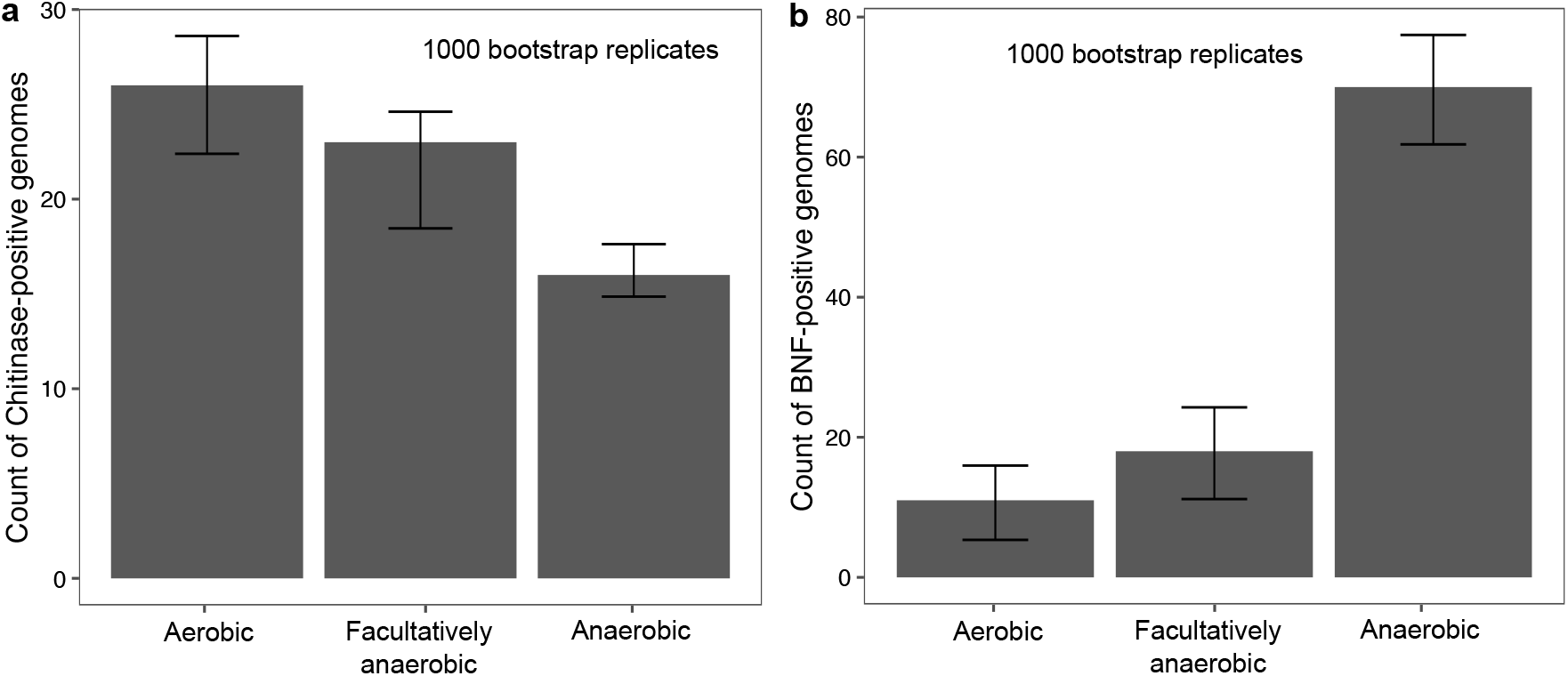
Lineages with chitinase genes are predominantly aerobic or facultatively aerobic while those with BNF potential are predominantly anaerobic. The number of (a) chitinase- and (b) BNF-positive genomes across aerobic, anaerobic, and facultatively anaerobic lineages in the filtered GEMs dataset. Error bars indicate 95% bootstrap confidence intervals around the median.

BNF, in contrast, is significantly negatively associated with aerobic respiration (glm model; p-value <2e-16; Fig. 4b) and positively with anerobic metabolism (glm; p-value = 0.0048; Fig. 4b). This aligns with the oxygen toxicity of BNF, and further suggests that the prevalence of N acquisition strategies is not simply regulated by metabolic energy yield. Another example for such control is the typical absence of nitrite assimilation pathways among *Desulfovibronales* as nitrite interferes with their energy generation pathway due to its structural similarity with sulfite (25).

### Effect of habitat type on the prevalence of N acquisition strategies

We expected to find a direct relationship between N strategy distribution and the environmental distribution of a taxon, under the assumption that the relative fluxes of bioavailable N compounds characteristically vary across environments, leading to differential adaptations among microbes inhabiting diverse habitats. For example, microbes living in association with a host (e.g., pathogens, parasites, symbionts, and gut commensals) tend to undergo genome reduction due to various evolutionary forces pushing towards minimizing functional redundancy and bolstering the host-resident (inter)dependency (26,27). If the resident microbe is being supplied with organic compounds by the host, independently acquiring N (particularly inorganic N) would be redundant and less efficient for the microbe. Therefore, we should expect to see genome reduction favoring the deletion of inorganic N acquisition pathways in host-associated microbes. Similarly, free-living taxa inhabiting nutrient-rich environments can potentially afford to eliminate inorganic N acquisition strategies. We examined evidence for these expected patterns in the GEMs dataset.

Indeed, taxa inhabiting host-associated environments were found to be relatively more enriched in organic N acquisition pathways than free-living taxa (Fig. 5a; # t = 25.039, df = 3986.6, p-value < 2.2e-16). This might reflect the greater availability of organic N compounds in the host environments, which makes inorganic N pathways redundant/futile. In addition, examining the relationship between inorganic N uptake and habitat type, we observed that inorganic N pathways are relatively less frequent among genomes obtained from high-nutrient environments such as bioreactor or solid/liquid waste environments (Fig. 5b; light blue and yellow points). Genomes assembled from built environments (predominantly city subway systems) frequently harbor chitin degradation potential, in addition to most LMW and inorganic N uptake strategies (Fig. 5b; darker blue points). In addition to subway systems, chitin degradation is relatively more common among genomes assembled from permafrost and plant rhizosphere systems (Fig. 5b, S3), which may reflect greater relative availability of HMW N compounds in these systems.

**Figure 5:**
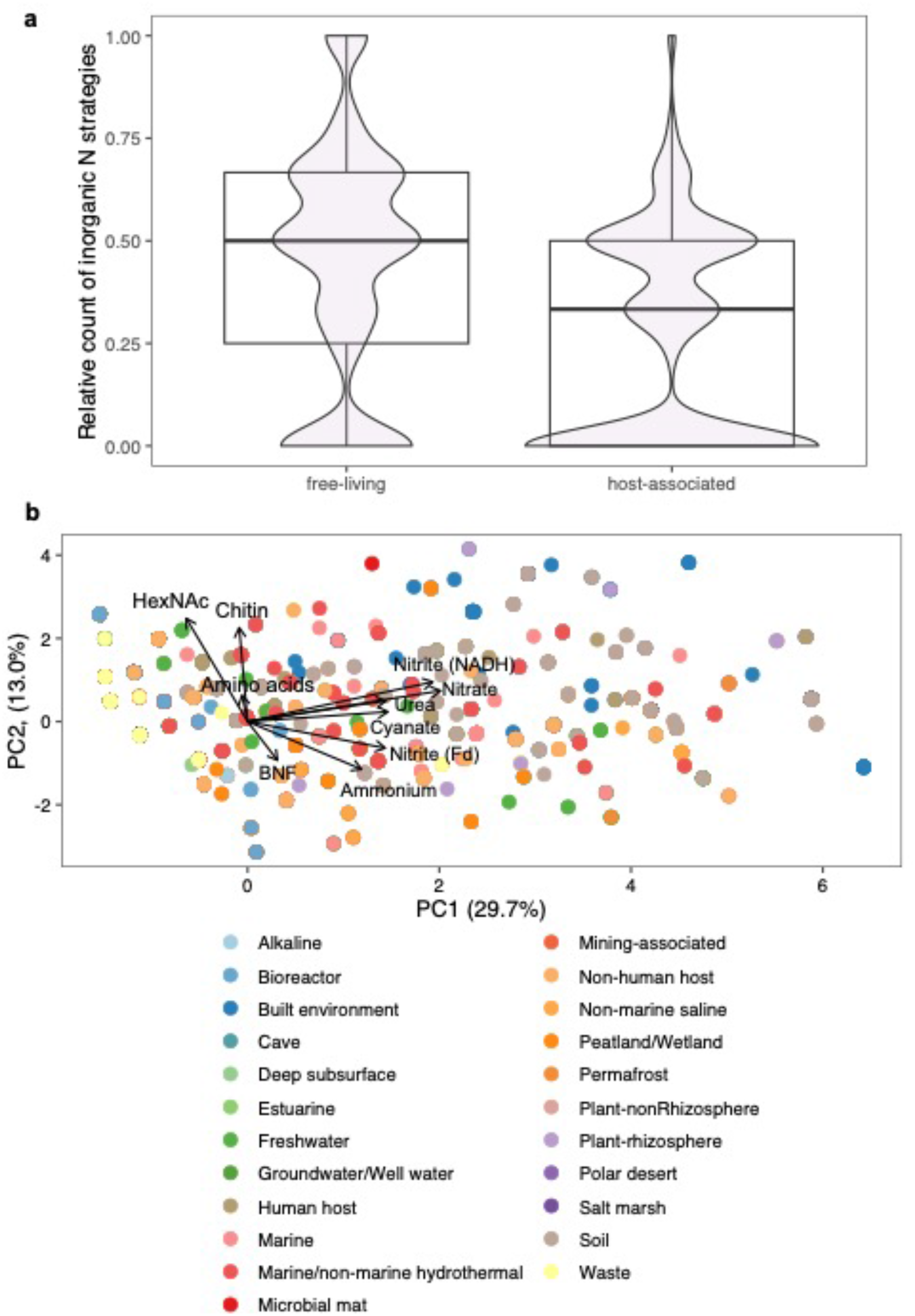
Relationships between environmental context and N acquisition strategy distributions. a. Number of inorganic-N acquisition strategies relative to the total number of N-acquisition strategies in free-living versus host-associated taxa. **b**. PCA analysis of the distribution of N acquisition strategies among the GEMs genomes across broad ecosystem categories.

We further tested associations between pathway incidence and ecosystem type using Poisson generalized mixed linear models with fixed effects for taxonomic relatedness at the family level. The results show significant associations between habitat type and N pathway distributions. There is a strong positive correlation between the number of organic N assimilation strategies and the following habitat types (corresponding p-values in parentheses): plant rhizosphere (<2e-16), built environment (i.e., city subways; <2e-16), human host-associated systems (1.9e-10), and freshwater systems (7.46e-14). Relatively weaker, yet significantly positive correlations were observed between the number of organic N strategies and aquatic marine systems (p-value 0.0002), soil (0.004), and solid/liquid waste environments (0.001).

These genomic observations most likely reflect differential availability of fixed N compounds across environments leading to microbial genomic adaptations to differences in N compound availability. For example, symbiotic taxa in host-associated environments may exhibit genome reduction, effectively lowering cellular energy requirements. This is observed in the case of the cyanobacterium *Candidatus Atelocyanobacterium* (UCYN-A), which is able to fix N_2_and supply fixed N to its host, but lacks any additional N-acquisition strategies, including ammonium uptake (28). Similarly, a large number of *Clostridia* genomes harbor chitinases and amino acid uptake systems yet lack additional pathways for N assimilation (including the typically prevalent ammonium uptake strategy; Table S2), and all of these turn out to be parasitic lineages that may be adapted to N-rich host environments.

An outstanding challenge in deciphering habitat-related patterns in pathway distributions is the lack of appropriate metadata for analyses. For example, environmental categories in the GEMs dataset are often too broad (e.g., “soil”, “freshwater”, or “marine”), which masks microhabitat diversity within each category. Environmental data at or close to the spatial scale of the microbe will be essential for making robust inferences on N strategy distributions. Substrate availability and transport limitations can vary greatly across habitat types, which will add to the metabolic and ecological complexity of the microbial inhabitants.

## CONCLUSIONS

Our examination of the distribution of N-acquisition strategies across microbial genomes reveal strong controls by the relative energetic costs of assimilation, which is influenced by specific metabolic adaptations to habitats and environmental conditions. Long standing energy cost framework explains the findings that assimilation of relatively cheap N compounds (ammonium and simple amino acids) dominates over all other strategies. Broader metabolic adaptations related to energy availability are also important: the higher free energy yield per mole of carbon from oxygen respiration likely affords the retention of larger, more versatile genomes in aerobic (and facultatively aerobic) organisms compared to anaerobic ones living at the bottom of the redox ladder (such as sulfate reducers and methanogens). Exceptions to this general pattern can result from additional controls including toxicity (i.e., pathway incompatibility) due to interactions between metabolites. For example, oxygen toxicity of BNF potentially explains the general prevalence of this strategy among obligate and/or facultative anaerobes. Similarly, the general absence of nitrite reduction in sulfate reducers may be explained by toxicity resulting from the structural resemblance between nitrite and sulfite. Additional habitat-related aspects such as host-associated versus free-living, oligotrophic versus copiotrophic systems can exert an additional, secondary control on the incidence of N-acquisition strategies. However, for robust testing of these hypotheses using genomic data, we need environmental data measured at the scale of the microbe.

Several additional N-acquisition pathways that we did not analyze in this study include extracellular peptidases and carbohydrate-active enzymes involved in the degradation of N-containing polymeric compounds other than chitin. Also, the presence of a gene/pathway complement in the genome does not necessarily indicate actual use of that pathway. For example, many organisms harboring the genomic potential for BNF have not been shown to grow diazotrophically (e.g., *Holophaga foetida, Thermodesulfovibrio* sp.). This suggests additional complexity in controls on the realized niches of microbes that ultimately determines N compound preferences in different ecosystems. The selection pressure on genomes can be highly variable across ecological contexts, which can contribute to differential genome evolution patterns in different habitats. Our analyses shed light on some of the resulting patterns in N strategy distributions, and guide future experiments to examine the relative substrate preferences in organisms harboring multiple N-acquisition strategies. Finally, we emphasize the importance of accurate and microbially relevant metadata for enabling genomic analysis that can allow predictive modeling of microbial ecology and evolution.

## ACKNOWLEDGMENTS

This study was supported by the Carbon Mitigation Initiative at the High Meadows Environmental Institute at Princeton University. The author(s) are pleased to acknowledge that the work reported on in this paper was substantially performed using the Princeton Research Computing resources at Princeton University which is consortium of groups led by the Princeton Institute for Computational Science and Engineering (PICSciE) and Office of Information Technology’s Research Computing.

## CONFLICT OF INTEREST DISCLOSURE

The authors declare no conflict of interest.

## Notes

### Competing Interest Statement

The authors have declared no competing interest.

