## Supplementary figures 1-3 for "A Genomic View of Environmental and Life History Controls on Microbial Nitrogen Acquisition Strategies"

*corresponding authors

### Supplementary Figures

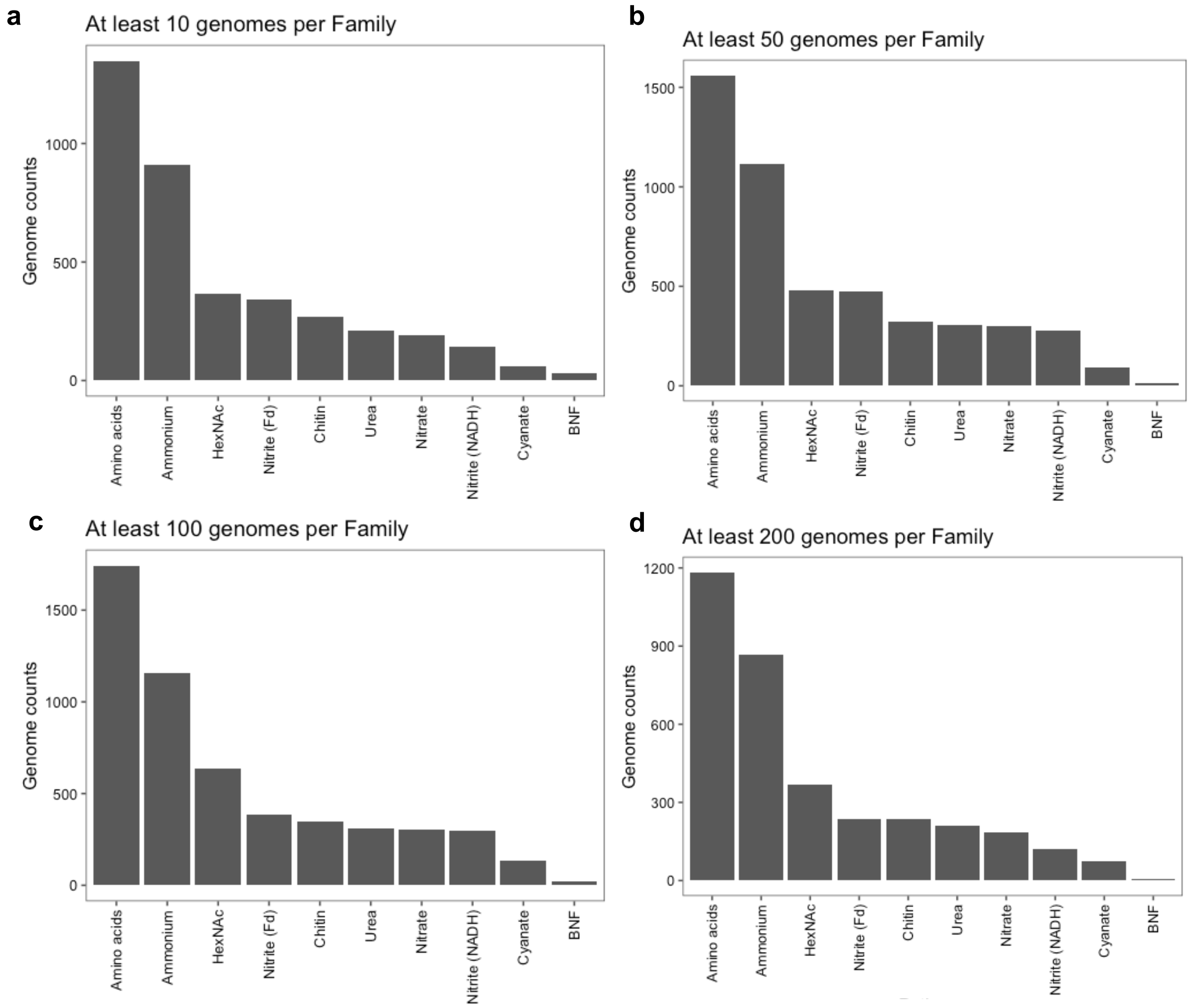


**Figure S1**: Prevalence of the various N-compound acquisition strategies among genomes in the filtered GEMs dataset, with different sample sizes. The dataset was filtered to retain family-level lineages that contain at least 10, 50, 100, or 200 genomes per family.

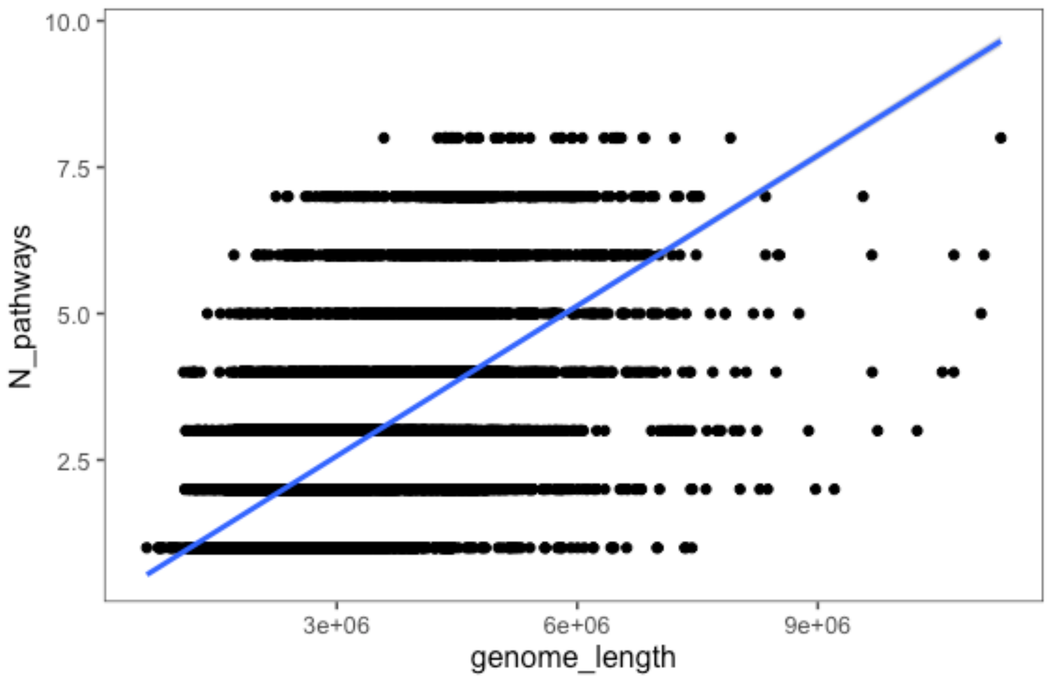


Figure S2: Correlation between genome length and the number of total N assimilation strategies.

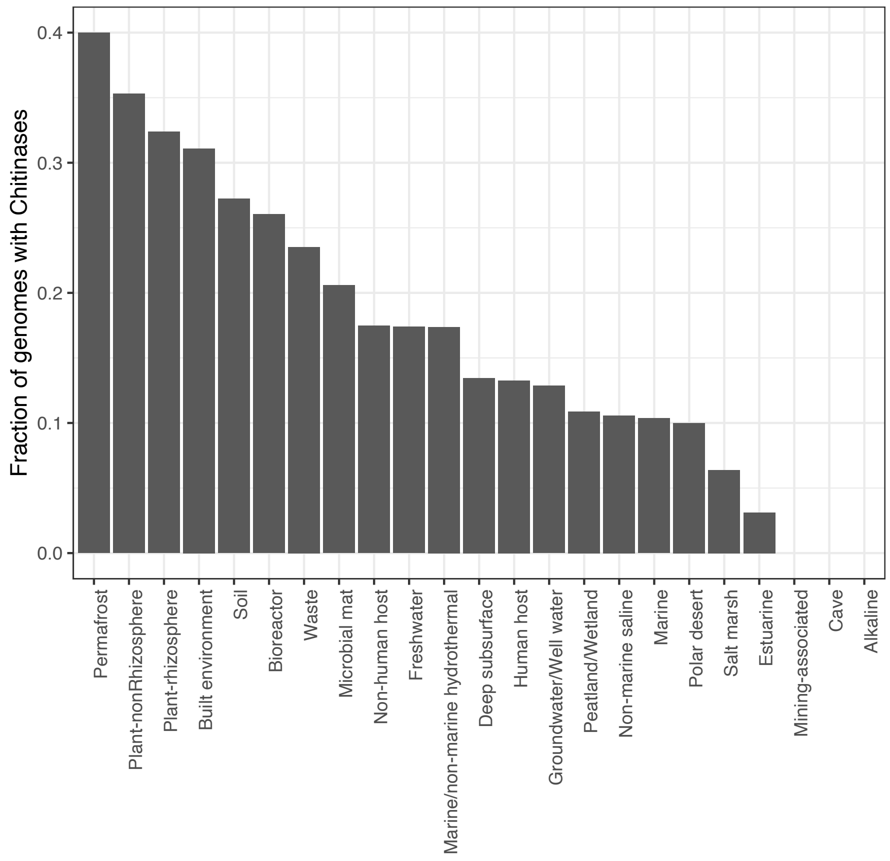


**Figure S3**: Relative counts of genomes with chitinases across habitat types.
